# Asparagine hydroxylation is likely to be a reversible post-translational modification

**DOI:** 10.1101/2020.03.22.002436

**Authors:** Javier Rodriguez, Cameron D Haydinger, Daniel J Peet, Lan K Nguyen, Alex von Kriegsheim

## Abstract

Amino acid hydroxylation is a common post-translational modification, which generally regulates protein interactions or adds a functional group that can be further modified. Such hydroxylation is currently considered irreversible, necessitating the degradation and re-synthesis of the entire protein to reset the modification. Here we present evidence that the cellular machinery can reverse FIH-mediated asparagine hydroxylation on intact proteins. These data suggest that asparagine hydroxylation is a flexible and dynamic post-translational modification akin to modifications involved in regulating signalling networks, such as phosphorylation, methylation and ubiquitylation.

## Introduction

Post-translational modifications (PTMs) are chemical alterations of amino acids or proteins which increase the complexity of the proteome and allow the cell to modify protein function in a dynamic or sustained manner (Ciechanover, 2005; Duan and Walther, 2015; Hunter, 2009; Walsh et al., 2005). Some of these modifications are irreversible, such as the cleavage of a polypeptide chain resulting in altered activity of the protein. Nevertheless, most PTMs are reversible and comprise the dynamic addition or elimination of functional groups that range from large oligosaccharide chains to a few atoms, such as glycosylation, phosphorylation, acetylation, methylation, or carboxylation. The reversibility of these PTMs is frequently achieved by the action of pairs of enzyme classes with opposing functions, one of which catalyses the forward reaction and another the reverse reaction (Gray and Teh, 2001; Tonks, 2006; Trewick et al., 2005). In cases where one of the reactions is thermodynamically unfavourable, the reaction may not be straightforwardly reversible and therefore includes intermediate products. An example would be the formation and dissolution of cysteine bonds, where the formation of the bond can include the oxidation of the sulphur of cysteine to sulfenic acid, which then forms a disulphide bond by reacting with another cysteine (Oka and Bulleid, 2013; Rehder and Borges, 2010)

Oxidation or more precisely hydroxylation of proteins on residues other than cysteine was recognised as a PTM in the 1960s, when the enzymatic hydroxylation of proline and lysine was identified as taking place during collagen synthesis (Hutton et al., 1967; Kivirikko and Prockop, 1967). Subsequently, the molecular mechanism of hydroxylation catalysed by the evolutionarily conserved family of the 2-Oxoglutarate dependent dioxygenases (2OG-ox) was described (Loenarz and Schofield, 2011; Ploumakis and Coleman, 2015). The 2OG-ox are a family of proteins composed of over 60 enzymes of which the so-called HIF-hydroxylases form a subgroup consisting of four hydroxylases: 3 prolyl hydroxylases (PHD1,2 and 3) and an asparaginyl hydroxylase “factor inhibiting HIF” (FIH). HIF-hydroxylases act as sensors of the oxygen levels within the cells. Under normoxic conditions prolyl hydroxylases (PHD1, 2 and 3) catalyse the specific hydroxylation of two proline residues in the alpha subunit of HIF1, the master regulator of the hypoxic response. Once hydroxylated, HIF1α is bound by the von Hippel-Lindau ubiquitin ligase (VHL) complex, which promotes the ubiquitination and rapid proteasomal degradation of HIF1α, resulting in the ablation of the protein under normoxic conditions (Appelhoff et al., 2004; Ivan et al., 2001; Jaakkola et al., 2001; Maxwell et al., 1999; Yu et al., 2001). FIH, the other component of the subfamily, catalyses an asparagine hydroxylation in the C-terminus of HIF1α. Upon hydroxylation of this residue, the interaction with co-factors required for the formation of the active transcription factor complex is impeded, resulting in the downregulation of HIF-driven transcription in the presence of oxygen (Lando et al., 2002b; Mahon et al., 2001).

Over the past few years, it has been argued that HIF1α is not the only protein that is hydroxylated by HIF-hydroxylases and several additional substrates, particularly of FIH, have been postulated and validated (Cockman et al., 2009a). In a similar manner to that observed in the context of HIFα, hydroxylation of these other substrates by FIH alters the physicochemical properties of the hydroxylated domains. These changes can induce or destroy protein-protein interactions that ultimately control substrate activity, folding or localisation (Janke et al., 2013; Karttunen et al., 2015; Kiriakidis et al., 2015; Rodriguez et al., 2016; Scholz et al., 2016).

Presently, it has been suggested that the hydroxylation mediated by HIF-hydroxylases is an irreversible process, and that the hydroxylation can only be reset by the degradation and new synthesis of a non-hydroxylated protein (Chan et al., 2005; Gorres and Raines, 2010; Schofield and Ratcliffe, 2004). A mass spectrometric study monitoring two FIH-mediated hydroxylation sites of Rabankyrin-5 substantiated this, as no evidence of dehydroxylation was found under the investigated conditions (Singleton et al., 2011). Nevertheless, some authors have proposed the existence of a dehydroxylation mechanism (Giaccia et al., 2003; Karttunen et al., 2015), but this notion is only founded on the assumption that PTMs should be generally reversible. Some data have emerged that implicitly suggest that asparagine hydroxylation may be a reversible PTM after all. First, in contrast to proline hydroxylation of HIF, FIH-mediated asparagine hydroxylation does not lead to the rapid decrease of protein half-life. This suggests that FIH-substrates are longer-lived proteins and that the cell would therefore benefit from a mechanism of resetting the level of hydroxylation in a more dynamic, non-destructive manner (Cockman et al., 2009a). One such example is transient receptor potential vanilloid 3 (TRPV3), an ion channel that is hydroxylated by FIH. Hydroxylation on an asparagine residue reduces TRPV3-mediated current, whereas hypoxia, FIH inhibition or mutation of the asparagine residue potentiates it without affecting protein stability. Intriguingly, the increases in current through the channel are observable in less than one hour of hypoxia or FIH inhibition. This rapid response indicates that there has to be a very rapid turnover of the protein or that the hydroxylation can be reversed without destruction and re-synthesis of TRPV3 (Karttunen et al., 2015). Moreover, indirect evidence from mathematical modelling indicates that signalling networks require reversibility of asparagine hydroxylation. Nguyen *et al* published a comprehensive ODE-based mathematical model of the immediate HIF network (Nguyen et al., 2013). Intriguingly, the mathematical model assumes irreversibility of both proline hydroxylations but includes an undefined reaction that leads to the dehydroxylation of the N-terminal asparagine residue (Fig. 1A). The reversibility of the asparagine hydroxylation was required for the model to reproduce the experimental data, which showed the transient induction of HIF protein levels and transcriptional output. Upon removal of the unspecified dehydroxylation reaction, the model predicts that HIF levels and activity would increase in a sustained manner, which is at odds with the experimental observations (Fig. 1B-E).

**Figure 1.**
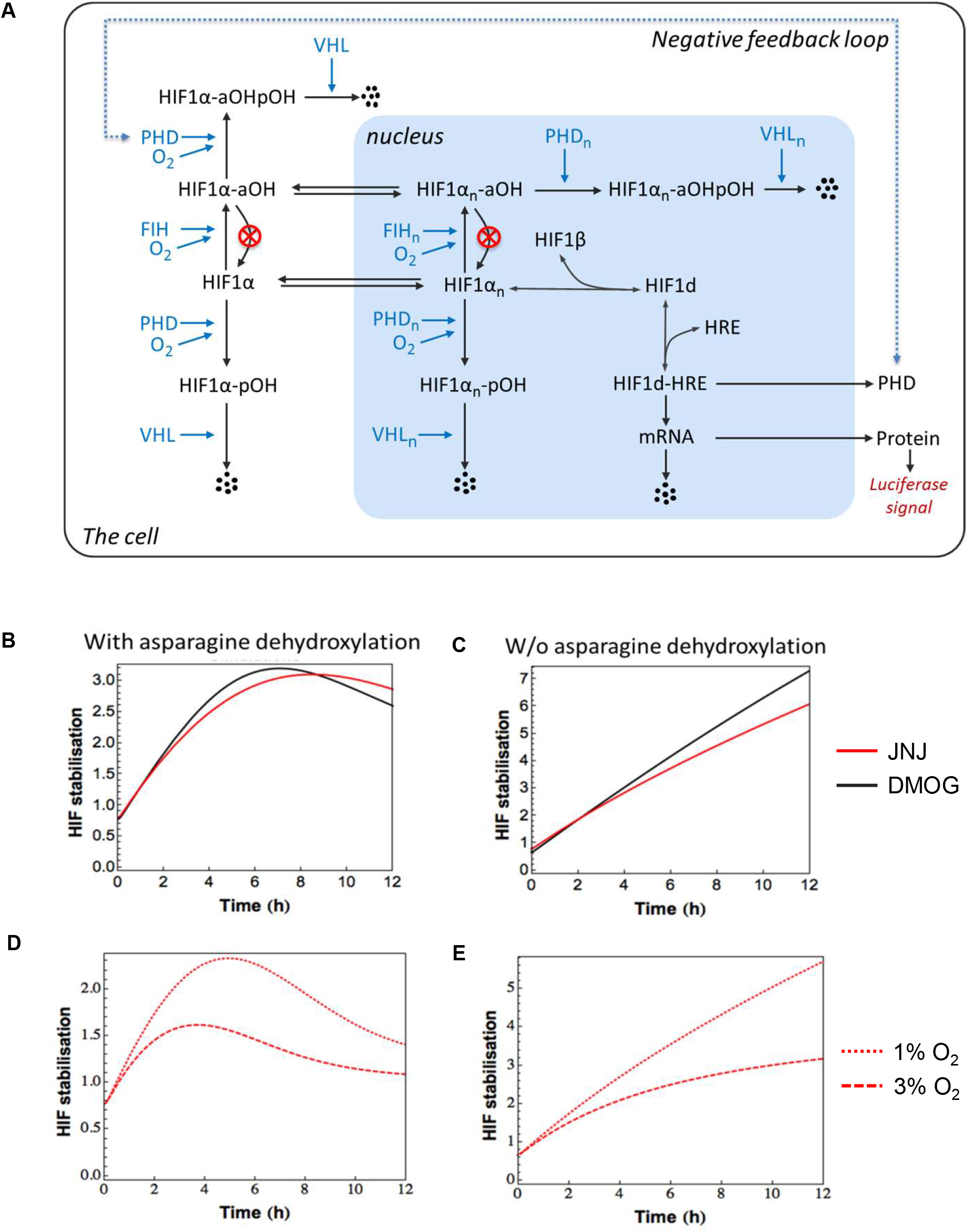
(A) Schematic diagram of a modified kinetic model of the HIF-1α signalling network where the assumed reactions representing asparagine dehydroxylation are removed from the original model (Nguyen et al., 2013) (denoted by the red crossed lines). (B-C) Comparison of the in silico predicted effects of DMOG or JNJ1935 on HIF-1a stabilisation in the presence (B) or absence (C) of asparagine dehydroxylation, the former match the transient dynamics observed in experimental data (Nguyen et al., 2013) while the latter display sustained patterns instead. (D-E) Comparison of the in silico predicted effects of 1% or 3% hypoxia on HIF-1a stabilisation in the presence (D) or absence (E) of asparagine dehydroxylation; the former match the transient dynamics observed in experimental data (Nguyen et al., 2013) while the latter display sustained patterns.

Overall, although these data give initial foundation to the hypothesis that asparagine hydroxylation is reversible, no direct evidence of reversibility has been produced. Notably, amino acid methylation was initially thought to be a static modification (Ramchandani et al., 1999), but was subsequently demonstrated to be reversed through the action of 2OG-ox. With this in mind, we set out to examine the potential reversibility of asparagine hydroxylation by using quantitative mass spectrometry.

## Results

There are several analytical techniques that can be applied to the quantitation of changes in PTMs, such as the use of modification-specific antibodies or the monitoring of shifts in the apparent molecular weight in PAGE (Luo et al., 2014; Snell et al., 2014). Unfortunately, neither of the above-mentioned methods is universal and for this reason, we decided to use quantitative mass spectrometry (qMS) to investigate whether hydroxylation is a reversible process. qMS has been shown to be superior to western blotting in terms of relative quantification and is a method widely used for monitoring changes in the hydroxylation status of proteins (Jaakkola et al., 2001; Luo et al., 2014; Rodriguez et al., 2016; Scholz et al., 2016; Singleton et al., 2011). Initially, we planned to monitor the dynamics of asparagine hydroxylation on the well-described FIH-substrate Tankyrase 2 (TNKS2).

Tankyrases are members of the poly(ADP-ribose)polymerase (PARP) protein super family, which participate in the regulation of the degradation complex in the Wnt/β-catenin signalling pathway (Mariotti et al., 2016; Waaler et al., 2012; Yang et al., 2016). TNKS2 has been characterized as a FIH substrate that contains the FIH consensus motif [LXXXXXV/IN] in several ankyrin repeat domains (ARD) (Cockman et al., 2009a; Cockman et al., 2009b). As such, TNKS2 contains multiple FIH-hydroxylation sites, and furthermore hydroxylation on TNKS2 by FIH has not been reported to have any effect on its stability. Multiple hydroxylations would allow us to monitor several sites in the same experiment, thus improving our chances of detecting if one or more hydroxylation(s) are reduced over time with statistical significance.

As an experimental approach we adapted the pulsed stable isotope labelling with amino acids in cell culture (SILAC), that has been successfully used to measure protein and PTM turnover (Boisvert et al., 2012; Ohsumi, 2006). HEK 293T cells were transfected with a V5-tagged FIH and a Flag-tagged TNKS2. 24 hours after transfection the medium was replaced with SILAC medium (containing ^13^C- and ^15^N-labelled arginine and ^13^C- and ^15^N-labelled lysine amino acids (R10K8)) and cell lysates were prepared at different time points. To further reduce the potential incorporation of unlabelled amino acids into proteins, we additionally worked in serum-free conditions, therefore, the comprehensiveness of labelled amino acid incorporation was only limited by the isotopic purity (generally >99%) and by the recycling of unlabelled amino acids resulting from the degradation of unlabelled proteins. Within these restrains, pulsing with heavy SILAC amino acids allowed us to distinguish TNKS2 synthesized post-pulse (Heavy SILAC) from the TNKS2 present before SILAC addition (light SILAC). To avoid possible rehydroxylation of peptides after the SILAC pulse, cells were treated with dimethyl-oxalylglycine (DMOG) and the lysis buffers contained N- oxalylglycine (NOG), both of which are pan-hydroxylase inhibitors (Bruick and McKnight, 2001). Therefore, if the hydroxylase inhibition is complete, heavy-SILAC peptides should be devoid of hydroxylation. By tracking how the ratio between hydroxylated and non-hydroxylated peptide intensity changes in the light SILAC TNKS2, we can monitor if a proportion of the hydroxylation is reversed (Fig. 2A). This experimental setup allows us to track the asparagine hydroxylation levels of the initial population of TNKS2 (light) and to determine how effective DMOG inhibited the hydroxylation reaction by assaying the relative hydroxylation changes in heavy-SILAC TNKS2. To establish the concentration required to inhibit FIH-mediated TNKS2 hydroxylation, we pulsed cells with increasing concentrations of DMOG and heavy SILAC media for 4 hours. We then quantified the heavy-labelled percentage of hydroxylated peptide vs. the vehicle control to assess the relationship between DMOG concentration and FIH inhibition. As expected, increasing concentrations of DMOG inhibited FIH more effectively (Fig. 2B, Fig S1A), with 2 mM reducing the activity by 98%. 1 mM and 0.5 mM DMOG only partially inhibiting FIH, consequently we used the highest concentrations for all further experiments

**Figure 2.**
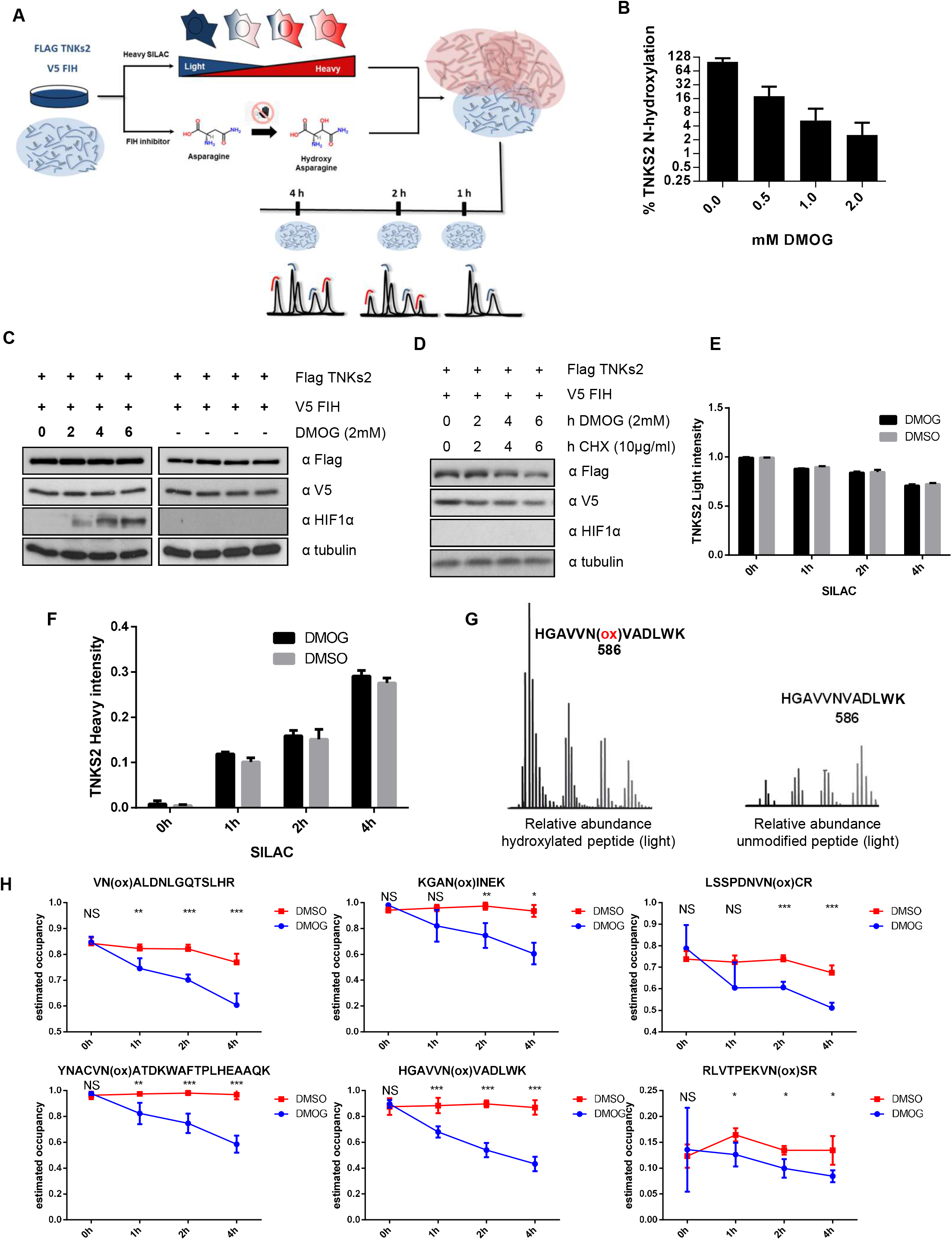
(A) Schematic illustration of the mass spectrometry based hydroxylase screen. (B) HEK293T cells were transfected with FLAG-tagged TNKS2. 24hours post-transfection the media was replaced by heavy SILAC and cells were treated with different concentrations of DMOG or vehicle for four hours. FLAG-tagged proteins were immunoprecipitated, digested and analysed by mass spectrometry. Graph represents the average percentage of heavy-labelled hydroxylated TNKS2 peptides in the presence of DMSO/DMOG, with 100% set to the DMSO level n=2, 6 peptides. (C) HEK293T cells were transfected with FLAG-tagged TNKS2. 24hours post-transfection the media was replaced by heavy SILAC and cells were treated with different times of DMSO or DMOG. The cells were lysed and proteins separated on PAGE, electroblotted and detected with the indicated antibodies. (D) HEK293T cells were transfected with FLAG tagged TNKS2 and V5 FIH. 24 hours after transfection cells were treated with CHX and DMOG. Cells were harvested and lysed at different times of CHX/DMOG/SILAC treatment, followed by western blot analysis with the indicated antibodies. (E) FLAG-tagged proteins were immunoprecipitated, digested and analysed by mass spectrometry. Diagram represent the intensities of immunoprecipitated light FLAG-TNKS2 across the DMSO/DMOG treatment. Error bars are standard deviation n=6. (F) As in E but representing heavy FLAG-TNKS2. (G) XIC of hydroxylated/unmodified light TNKS2 peptide (HGAVVNVADLWK) along DMOG treatment from 0, 1, 2 and 4 hours. (H) Graphs represent the estimated hydroxylation occupancy of light SILAC TNKS2 peptides. The error bars represent standard deviation and n = 6.

To determine whether SILAC media and DMOG affected TNKS2 protein stability, we first validated by western blotting that the expression of TNKS2 or FIH was not affected by either treatment at any time point (Fig. 2C). Furthermore, to estimate TNKS2 protein turnover we blocked protein synthesis with cycloheximide, a rapidly acting eukaryotic protein synthesis inhibitor (Kay and Korner, 1966; McMahon, 1975) commonly used for protein turnover studies (Larance et al., 2013). We initially monitored how CHX inhibited protein synthesis by western blotting and observed a reduction in the levels of Flag-TNKS2 and V5-FIH, with an approximate half-life of 6 hours for TNKS2 (Fig. 2D). We next determined how TNKS2 hydroxylation levels change dynamically upon SILAC and DMOG pulsing. We immunoprecipitated Flag TNKS2, digested the protein on-beads, analysed the peptides by LC-MS/MS and quantified TNKS2 expression using MaxQuant. This confirmed our initial observation that DMOG did not affect the stability of TNKs when compared to the DMSO control (Fig. 2E) and also allowed us to monitor the incorporation of heavy SILAC, by detection of the increase in heavy-labelled TNKS2 (Fig. 2F). Based on the labelling data, we calculated the TNKS2 protein half-life to be around 7 hours. This matched the Western-blot-based estimation, suggesting that newly synthesised TNKS2 protein is predominantly heavy-labelled upon pulsing with heavy amino acids and that DMOG treatment does not affect the expression and stability of exogenously expressed TNKS2.

Having confirmed this, we began using this set-up to determine if hydroxylated proteins could be dehydroxylated in cells. Initially, we analysed the ion chromatogram of the light-labelled TNKS2 peptides containing hydroxylated and non-hydroxylated asparagine 586, and detected a reduction in the intensity of the hydroxylated peptide associated with DMOG treatment, whilst at the same time we detected an increase of the absolute levels of the unmodified peptide (Fig. 2G), suggesting that hydroxylated residue can be reverted. To increase certainty, we increased the number of experimental repeats to six and analysed several hydroxylation sites in an automated manner. We selected the peptides that we identified with a localization-specificity for hydroxylated asparagine of greater than 0.66 and where both the hydroxylated and non-hydroxylates isoform were detected in at least four of the six replicates. The hydroxylation occupancy of the peptide was estimated by transforming the ratio of the hydroxylated over non-hydroxylated peptide (determined by the search software MaxQuant) using following formula: 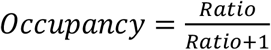 This transformation assumes that the molar ionisation efficiencies of both peptide species are equivalent. Generally speaking, this is plausible as the physico-chemical properties of the peptides are similar, exemplified by a small shift in retention time.

Using this automated analysis method we monitored how the occupancy of several light-labelled hydroxylated peptides changed over time (Fig. 2H). Six sites showed a statistical significant decrease in hydroxylation occupancy relative to a negative control. The occupancy of some sites was only altered by a few percentage points, whereas others by up to 50%, suggesting a gradient of reversibility.

To rule out the possibility that the pulsing with heavy amino acid did not comprehensively abrogate protein synthesis of light-labelled protein, we repeated the experiment using CHX as an additional control. We combined CHX with a heavy SILAC and DMOG/DMSO pulse in order to abrogate TNKS2 synthesis during the time course of DMOG treatment. Heavy-SILAC pulsing further allowed us to determine how efficiently CHX abrogated protein synthesis. We initially monitored how CHX inhibited protein synthesis by LC-MS and observed a near complete abrogation of heavy-labelled TNKS2 production when compared to a control (Fig. 3A). Overall, TNKS2 synthesis was inhibited by over 98% in the presence of CHX and can therefore be considered residual. Once this was confirmed, we immunoprecipitated Flag TNKS2 from HEK293T cells post pulse and analysed the hydroxylated by LC-MS/MS as above. When monitoring how the relative occupancy of the light-labelled hydroxylation changed upon pulsing with DMOG, we were able to observe a significant reduction of several asparagine residues (Fig. 3B-D). Surprisingly, the reduction of the hydroxylation occupancy observed was smaller when compared to the previous experiments. We further failed to observe the proportionality between incubations times and the reduction in hydroxylation occupancies. This initially puzzling observation could suggest that what we overestimated dehydroxylation in the previous experiment due to a significant *de novo* synthesis of light-labelled TNKS2. If this would be the case, roughly three quarters of the newly synthesised TNKS2 would have to be light labelled. This is inconsistent with our previous observations where half-life estimations based on heavy-label incorporation match estimations based on CHX-pulse experiments. An alternative explanation is that the dehyxdroxylase or an essential co-factor is rapidly degraded upon CHX treatment. Should the enzyme/s catalysing the dehydroxylation reaction have a short half-life, this would result in partial dehydroxylation during the initial time points of the experiment, followed by a plateau. This is a profile not dissimilar to what we have observed (Fig. 3B-D).

**Figure 3.**
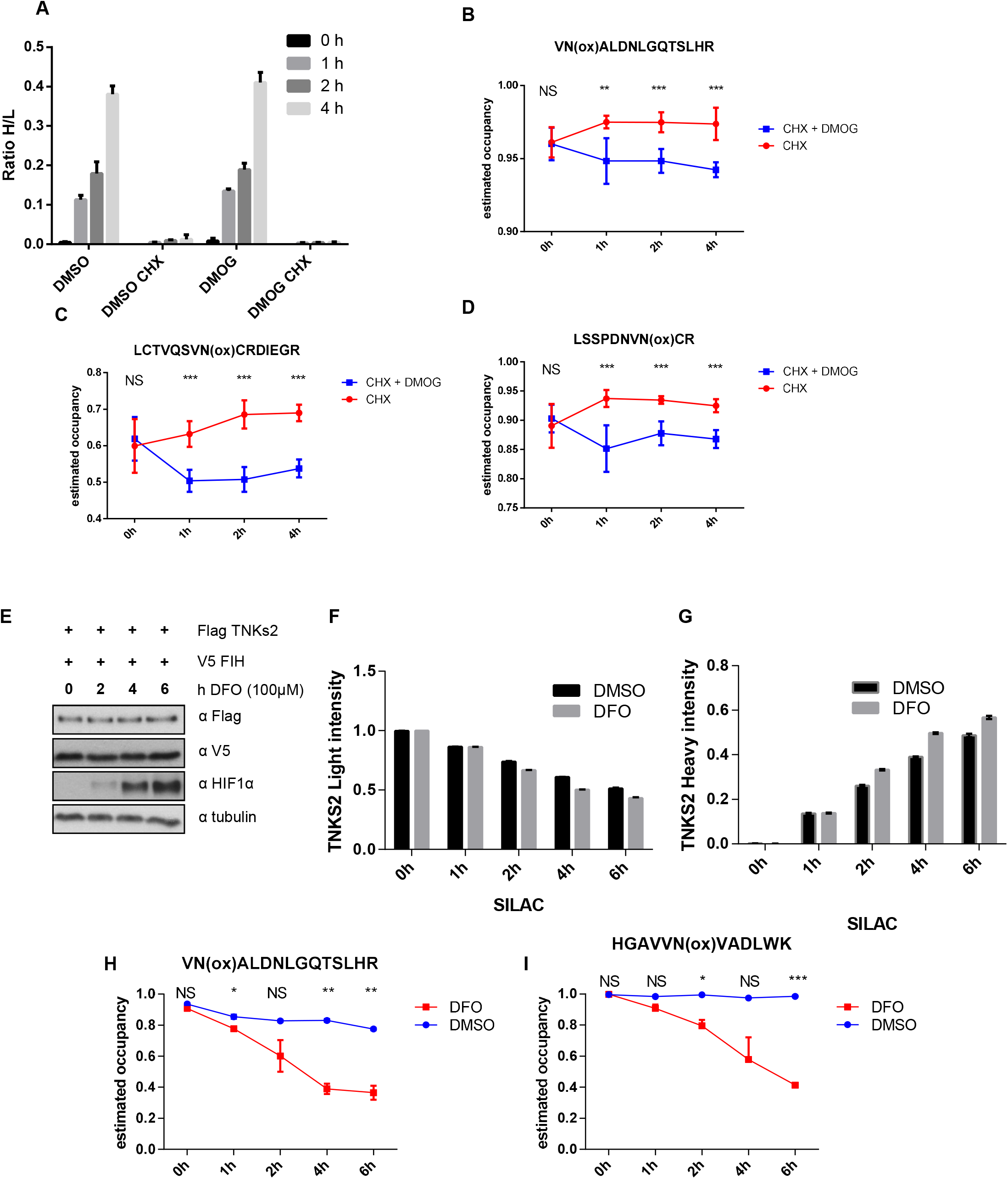
(A) HEK293T cells were transfected with FLAG tagged TNKS2 and V5 FIH. 24 hours after transfection cells were treated with CHX and DMOG and the media was replaced by heavy SILAC. Bar graphs represent the ratio of intensity heavy/light SILAC of FLAG TNKS2 during the control/CHX treatment. The error bars represent standard deviation and n = 6 (B-D) Graphs represent the estimated hydroxylation occupancy of light SILAC TNKS2 peptides. The error bars represent standard deviation and n = 6. (E) HEK293T cells were transfected with FLAG tagged TNKS2 and V5 FIH, the media was replaced by heavy SILAC and cells were treated with DFO. Cells were harvested and lysed at different times of DFO/SILAC treatment. Proteins were separated by PAGE, electroblotted and detected with the indicated antibodies. (F) FLAG-tagged TNKS2 proteins were immunoprecipitated, digested and analysed by mass spectrometry. Bar graphs represent the intensities of immunoprecipitated light FLAG-TNKS2 across the DFO treatment. Error bars are standard deviation n=2. (G) Bar graphs represents the intensity of heavy FLAG TNKS2 during the DFO treatment. Error bars are standard deviation n=2. (H-I) Graphs represent the estimated hydroxylation occupancy of light SILAC TNKS2 peptides. The error bars represent standard deviation and n = 2

To rule out a non-specific side-effect of DMOG treatment, we utilised a second, structurally and functionally unrelated inhibitor, deferroxamine (DFO), which inhibits FIH by chelating the iron in the active centre (Prass et al., 2002; Sharp and Bernaudin, 2004). Firstly, as we did previously for DMOG, we checked by western blot and mass spectrometry that DFO did not affect the stability of the protein (Fig. 3E) and the incorporation of heavy SILAC (Fig. 3F, 3G). Secondly, we quantified the occupancy for two previously characterised sites. As previously, we observed a reduction in the hydroxylated/non-hydroxylated ratio upon DFO treatment in the Light SILAC samples (Fig. 3H, 3I).

To determine whether dehydroxylation takes place on other substrates we decided to investigate the reversibility of a hydroxylation site on TRPV3. As mentioned in the introduction, TRPV3 is hydroxylated by FIH on asparagine 242 (Karttunen et al., 2015) and the hydroxylation regulates TRPV3 activity without affecting its expression. Interestingly, TRPV3 activity responds rapidly to hydroxylase inhibition, suggesting that dehydroxylation could be taking place. Replicating our experimental setup in HEK 293T cells and over-expressing myc-tagged TRPV3, we initially confirmed that inhibition of FIH by DMOG and incubation with SILAC media did not affect TRPV3 protein expression. By western blotting we confirmed previous observations that TRPV3 protein stability was not affected by hydroxylase inhibition (Fig. 4A). We then immunoprecipitated myc-TRPV3, digested with trypsin and analysed resulting peptides by mass spectrometry. Overall intensity of the light-labelled TRPV3 was altered over 2 hours and we detected less than 20% incorporation of the heavy label (Fig. 4B), suggesting that TRPV3 is a protein with a long half-life. In addition, we readily detected the reported hydroxylation of N242. We could further determine that for light-labelled N242 the hydroxylation occupancy was rapidly reduced over time upon DMOG treatment (Fig. 4C). Together, these data suggest that the hydroxylation of N242 in TRPV3 is reversible.

**Figure 4.**
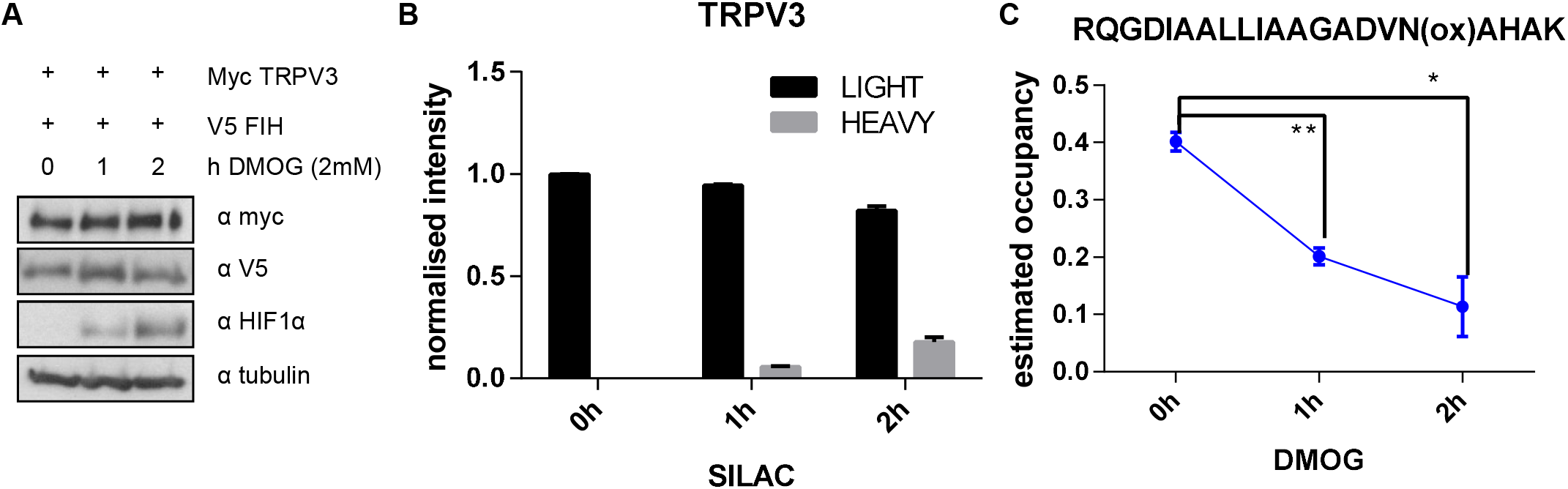
(A) HEK293T cells were transfected with myc tagged TRPV3 and V5 FIH. 24 hours after transfection the media was replaced by heavy SILAC and cells were treated with DMOG and. Cells were lysed at different times of DMOG/SILAC treatment. The expression of the indicated proteins were analysed by western blot (B) Myc-tagged proteins were immunoprecipitated, digested with trypsin and analysed by mass spectrometry. Diagram shows the intensities of light and heavy myc-TRPV3 during the DMOG treatment. Error bars are standard deviation n=2. (C) Graph shows the estimated hydroxylation occupancy of light SILAC TRPV3 peptide during DMOG treatment. Error bars show standard deviation and n = 2.

Finally, we checked if our finding could be extrapolated to the best-studied substrate of FIH, the alpha subunit of HIF1α (Lando et al., 2002b; Mahon et al., 2001). HEK293T were transfected with a GFP tagged HIF1α plasmid in order to overexpress the protein. As previously, we checked if the total levels of GFP HIF1α were affected by DMOG or heavy SILAC treatments. By western blotting we surprisingly detected that HIF1α expression appeared to be stable under normoxic conditions and that DMOG only marginally, if at all, increased HIF1α protein levels (Fig. 5A). We repeated the same experiment, and using mass spectrometry we observed a reduction in the levels of light-labelled HIF1α, at the same time that we detected an induction of the heavy-labelled GFP HIF1α (Fig. 5B). Finally, we studied the oxidation levels of N802 (the asparagine residue hydroxylated by FIH) and we observed that the ratio of oxidation of N802 was rapidly reduced upon DMOG treatment (Fig. 5C). As we expressed full-length HIF1α we decided to additionally monitor the hydroxylation of both reported proline sites. Whereas we identified the unmodified peptides with ease and over 50 MS/MS each (data not shown), we were not able to detect a single MS/MS identifying the proline hydroxylation with confidence. We have therefore concluded that overexpressing HIF1α overwhelms the capacity of the endogenous PHD1−3 enzymes to hydroxylate the proline residues stoichiometrically, delivering an explanation as to why we can detect HIF protein expression in normoxia. Overall, these results match the data that we obtained for TNKS2 and TRPV3, suggesting the presence of an asparagine dehydroxylation reaction. To determine whether other modifications can occur on the hydroxylated peptide we repeated the data analysis including the dependent-peptide matching option, which allows for the unbiased identification of modified derivatives of “base” peptides. The algorithm detected several modifications localised to N803, one of which was the elimination of two hydrogens (Fig. S2). We could only detect this modification on light-labelled peptides, suggesting that presence of DMOG reduces the amount of the precursor, indicating that didehydrogenated N803 is derived from hydroxylated N803. This would be consistent with an intermediate product in the potential dehydroxylation reaction where water is eliminated from the hydroxylated side-chain. Nevertheless, this is speculation as the abundance of didehydrogenated peptide is miniscule preventing us from reliably quantifying it across the samples.

**Figure 5.**
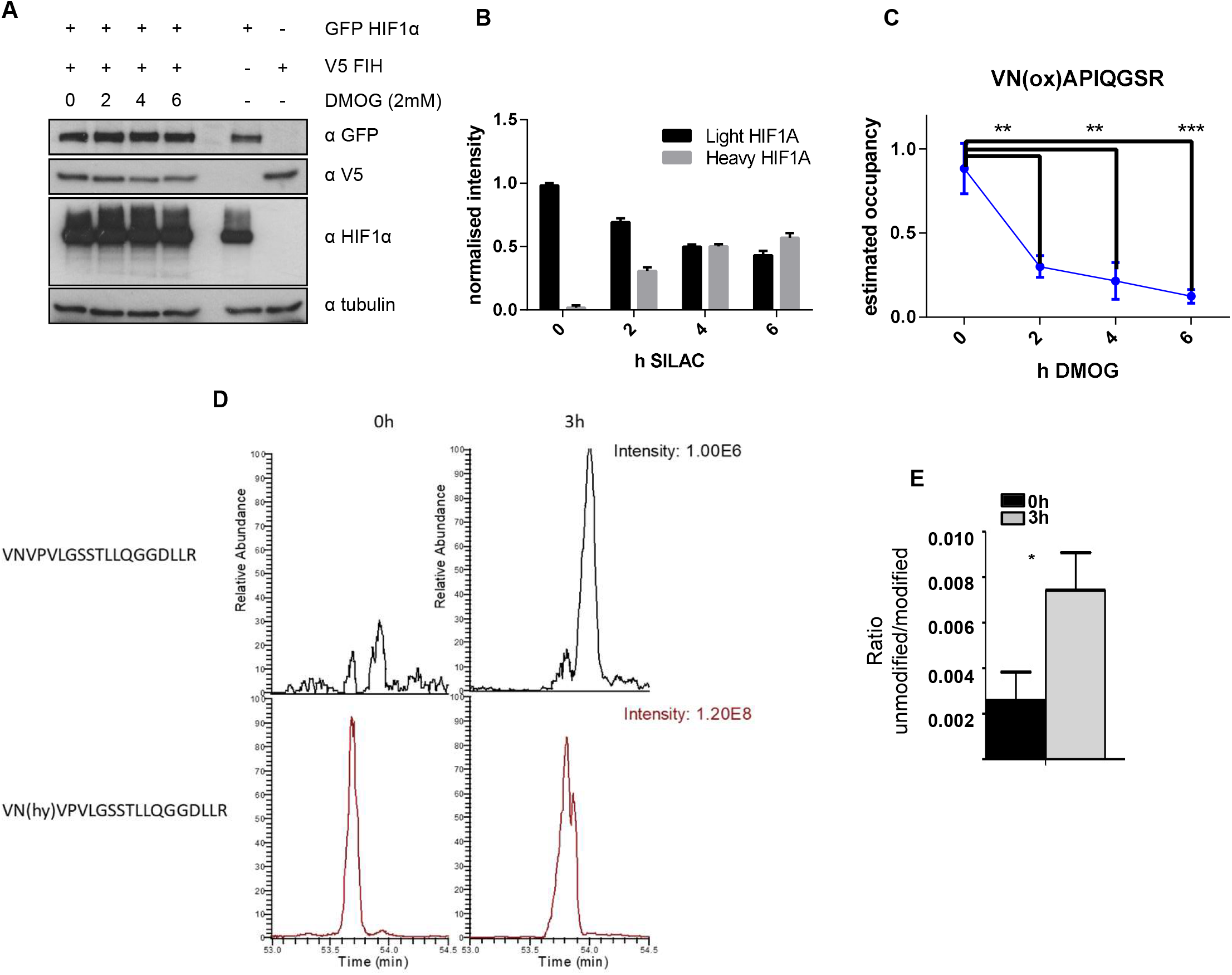
(A) HEK293T cells were transfected with GFP HIF1α and V5 FIH. 24 hours after transfection cells were treated with DMOG and the media was replaced by heavy SILAC. Cells were harvested and lysed at different times of DMOG/SILAC treatment. Total lysates were probed for the indicated proteins for immunoblotting. (B) GFP-tagged proteins were immunoprecipitated, digested with Trypsin and GluC and finally analysed by mass spectrometry. Bar graphs represent the intensities of immunoprecipitated light and heavy GFP-HIF along the DMOG treatment. Error bars are standard deviation n=2. (C) Graph shows the estimated hydroxylation occupancy of light SILAC HIF peptide during DMOG treatment. Error bars represent standard deviation and n = 2. (D) Representative XIC of unmodified and hydroxylated HIF2α peptide before and after incubation for 3 hours with FIH-/-HeLa lysate. (E) Bar graph shows the hydroxylation ratio of the unmodified over hydroxylated HIF2α peptide before and after incubation with FIH-/-HeLa lysate. Error bars represent SEM and n = 3.

To determine if we could observe asparagine dehydroxylation in a cell-free reaction, we generated hydroxylated protein by incubating the recombinant C-terminal domain of HIF2α (CAD) with recombinant FIH. To prevent re-hydroxylation by FIH, hydroxylated CAD was re-purified and subsequently incubated with cellular lysates derived from FIH-knockout HeLa cells (Chen et al., 2017). After 3 hours the CAD was affinity purified, digested and analysed using mass spectrometry. The ratio of non-hydroxylated peptide over hydroxylated peptide was approximately double that of the control sample that was immediately purified upon mixing with the lysate (Fig. 5D-E). Importantly, we observed an increase in non-hydroxylated peptide. Although the overall amount is small, the ratio increased to less than 1/100 of the hydroxylated intensity, this is not unexpected given the supraphysiological levels of hydroxylated peptide that were incubated with the HeLa cellular lysates.

Taking together, the data obtained from three different FIH substrates supports the hypothesis that asparagine hydroxylation is a post-translational modification that can be reversed within cells, with additional evidence that the hydroxylation can be reversed *in vitro*.

## Discussion

Protein functions can be switched on and off by distinct PTMs, such as phosphorylation, glycosylation and others (Ciechanover, 2005; Duan and Walther, 2015; Hunter, 2009; Walsh et al., 2005), which allows cells and organisms to respond dynamically to changes in the environment. The postulated reversibility of hydroxylation could explain how acute hypoxia and reoxygenation elicits rapid responses irrespective of protein degradation. This may be especially important in tumours as cyclical/intermittent hypoxia and protein hydroxylation are cancer hallmarks (Araos et al., 2018; Ruan et al., 2009).

We have not yet identified the enzyme/s responsible for the postulated reaction. We initially took a candidate approach and knocked-down proteins that were commonly binding to several N-hydroxylated proteins. However, none of these pertrubations had an effect on the base-line hydroxylation levels (data not shown), which suggests that the best option would be to devise a genome-wide screen and using N-hydroxylated-dependent protein-protein interactions or antibodies as a readout. Identification of the enzyme responsible would also allow design of a tailored *in vitro* assay that could reveal parameters and molecular mechanisms of the reaction, the precise nature of which we can currently only speculate on. Unexpectedly, the experiments we conducted in presence of CHX may have narrowed the search area. It is attractive to speculate that the dehydroxylase is a short-lived protein, as this would enable rapid and dynamic regulation of its activity. Rapid degradation would also bestow the system with the ability to adapt to the cellular needs dynamically. It is also plausible that such a system could include additional regulation downstream of hydroxylases, either as transcriptional target of the HIF-pathway or as direct PHD/FIH substrate. Indeed, most reported PHD-mediated hydroxylations result in protein stabilisation, although some discussion has emerged whether PHDs have any substrates apart from HIFs (Cockman et al., 2019). Either way, such feedforward loops are extensively used in signalling networks as they increase the dynamic response of a system when facing perturbations. Furthermore, rapid turnover of the postulated dehydroxylase delivers a possible explanation as to why dehydroxylation was not observed using CHX-chase experiments (Singleton et al., 2011). A further difference between our work and attempts to detect dehydorxylation, is the addition of NOG to our buffers. This prevents re-hydroxylation post-lysis when the concentration of the intercellular inhibitor are diluted, possibly a crucial step if longer processing of non-denatured samples is required, such as during immunoprecipitations.

When searching for the potential dehydroxylase we can also look at how similar reactions are catalysed in cells. Typically, these involve the formation of a short-lived intermediate by elimination of water, followed by the hydrogenation of the resulting unsaturated bond. Unbiased screening of asparagine modifications indeed detected this potential intermediate, but future work will have determine whether this is the dehydroxylation mechanism. The discovery of the dehydroxylation enzyme/s would also provide a set of likely new therapeutic targets to treat cancer or other human pathologies. For this reason, identification of the enzyme should be a priority and is part of our plans.

## Materials and Methods

### Cell culture

HEK293T cells were cultured in Dulbecco’s modified Eagle medium (DMEM) supplemented with 2 mM glutamine (Invitrogen) and 10% foetal calf serum (Invitrogen). Plasmids were transfected with Lipofectamine 2000 (Invitrogen) according to the vendor’s instructions. SILAC media was generated by custom made DMEM medium lacking L-arginine and L-lysine (Thermo Scientific). This media was supplemented L-Arginine-^13^C ^15^N (R10) and L-Lysine-^13^C ^15^N (K8) (Cambridge Isotope Laboratories, Inc.) but not with FCS to prevent incorporation of ^12^C-Lysine or Arginine.

### Materials

All antibodies were from commercial sources: anti-FLAG M2 peroxidase was obtained from Sigma Aldrich (F4042, 1:1,000 dilution), anti-HIF1α was from BD Biosciences (610958 1:1,000 dilution), anti-GFP and anti-myc were from Cell signalling (D5.1-9B11, 1:2,000 dilution,) anti-tubulin was purchased from Santa Cruz (sc-8035, 1:1,000 dilution) and anti-V5 was obtained from Invitrogen (R96025, 1:5000 dilution). DMOG was obtained from Cayman Chemical (71210), DFO and cycloheximide were purchased from SIGMA (D9533-C4859). For immunoprecipitation anti-Flag-M2 beads (Sigma Aldrich), anti-myc beads (Cell signalling-9B11) and anti-GFP (GFP-Trap Magnetic Agarose-Chromotek) were used.

### Plasmids

Myc-TRPV3 was a gift from Dan Peet (University of Adelade), GFP-HIF1A was a gift from Alex Chong (Aston University), FLAG-TNKS2 from Addgene (#34691)

### Cell lysis and immunoblotting

Cells were lysed in ice-cold lysis buffer (1% Triton-x100, 20 mM Tris-HCl (pH 7.5), 150 mM NaCl), supplemented with protease (5 μg/ml leupeptin, 2,2 μg/ml aprotinin), phosphatase inhibitors (20 mM β-glycerophosphate). Lysates were cleared of debris by centrifugation at 20,000 x g for 10 min in a benchtop centrifuge (4°C). Total lysates were fractionated by SDS-PAGE and transferred onto nitrocellulose filters. Immuno-complexes were visualized by enhanced chemiluminescence detection (GE Healthcare) with horseradish peroxidase–conjugated secondary antibodies (Bio-Rad Laboratories).

### *In vitro* dehydroxylation reaction

Thioredoxin-6 histidine (Trx-6H) tagged human HIF-2α CAD (774-874) and maltose binding protein (MBP) tagged human FIH were expressed in BL21(DE3) E. coli as previously described (Lando et al., 2002a). Protein expression was induced with 0.5 mM IPTG. CAD was induced for 8 h at 30°C and purified using a HisTrap HP column (GE Healthcare). FIH was induced for 16 h at 16°C and purified using a MPBTrap HP column (GE Healthcare). Proteins were exchanged into 150 mM NaCl, 20 mM Tris-HCl pH 8 using PD-10 desalting columns (GE Healthcare). 10 μM CAD was hydroxylated by incubation with 10 μM FIH, 112.5 mM NaCl, 65 mM Tris-HCl, 4 mM sodium ascorbate, 500 μM DTT, 30 μM FeSO4 and 40 μM 2-oxoglutarate for 2.5 h at 37°C, and repurified using a HisTrap HP column (GE Healthcare). FIH -/- HeLa cells (Chen et al., 2017) were lysed in ice-cold lysis buffer (1% Triton-x100, 20 mM Tris-HCl (pH 7.5), 150 mM NaCl), supplemented with HALT EDTA-free protease inhibitors). Protein concentration was ~0.5mg/ml. 1 μg of hydroxylated CAD was incubated under shaking at 37°C with 1 ml of FIH -/- HeLa lysate for 3 hours (or 0 hours for control). Precipitated with Ni-NTA-agarose beads (Quiagen) and digested with Trypsin and GluC and processed as previously described (Turriziani et al., 2014). Mass spectrometry was done using a Lumos Fusion (Thermo) mass spectrometer coupled to a RLS-nano uHPLC (Thermo). Peptides were separated by a 40 minute linear gradient from 5-30% Acetonitrile, 0.05% Acetic acid. Mass range was 646.5-653.5. XIC were generated with a width of (646.6980-646.7140)/(652.0300-653.0440) for the for non-hydroxylated/hydroxylated peptide. Peptide elution times were calibrated using hydroxylated/non-hydroxylated standards.

### Mass spectrometry

HEK293T cells were transfected with the different plasmids indicated. The different treatments were done 24 hours post-transfection. The cells were lysed and we immunoprecipitated the protein transfected with anti-FLAG, anti-myc or anti GFP agarose for 2 hours. Lysis and wash buffers was supplemented with 1 mM NOG. All the samples were digested with trypsin and also with GluC for the GFP-HIF1α and processed as previously described (Turriziani et al., 2014).

### Data analysis

The mass spectrometry raw data was analysed by the MaxQuant software packages using the pre-selected conditions for LFQ analysis. Specifically, MS/MS spectra were searched against the human Uniprot database with a mass accuracy of 4.5 ppm and 20 ppm (for MS and MS/MS). Carbamylation (c) was selected as fixed modification. Variable modifications were N-terminal acetylation (protein) and oxidation (M) for the interaction expression screen, oxidation (MKPNH) for the hydroxylation screen. FDR was set to 0.01. Peak matching was selected and was limited to within a 0.7 min. elution window with a mass accuracy of 4.5 ppm. Normalisation of hydroxylated peptides was performed by dividing the intensity of the modified by the matching unmodified peptide. Ratio were converted into estimated occupancies by dividing the ratio by itself +1; (x/(x+1)); using the Perseus software. For the dependent peptide search MaxQuant (1.6.7) was used with the pre-configured parameters.

### Statistical analysis

Technical replicates were averaged. Protein intensities and hydroxylation occupancies are shown with error bars representing standard deviation.

### In silico modelling

To in silico predict the effects that putative dehydroxylation may have on dynamic behaviour of the HIF-1α signalling pathway, we modified a well-calibrated mathematical model of the HIF pathway which we developed previously (Nguyen et al., 2013), by removing the steps representing asparagine dehydroxylation in this model (Fig.1A). This was done by setting the kinetic parameters describing the rate of these reactions to null in the model’s ordinary differential equations. Simulations of the intact and the adjusted models under various conditions (i.e. hydroxylase inhibition by DMOG and JNK, and 1% and 3% hypoxia, Fig. 1B-E) show that removal of the asparagine dehydroxylation steps failed to reproduce the experimental patterns of HIF-1α expression.

### Data availability

MS-data is deposited at Proteomexchange PXD013116

## Acknowledgements

We thank Dan Peet and Alex Cheong for the TRPV and HIF1α plasmid; Andrew Finch and Arkadiusz Welman for corrections and critical reading, Cancer Research UK. Cancer Research UK (CRUK Edinburgh Centre C157/A255140), Wellcome Trust (Multiuser Equipment Grant, 208402/Z/17/Z) for funding.

## Author Contributions

J.R and AvK designed the experiments, interpreted the results, J.R. D.J.P and C.D.H performed most experiments. L.N. designed and interpreted the mathematical modelling, AvK, J.R, L.N. and D.J.P wrote and edited the manuscript.

---

**Figure S1.**
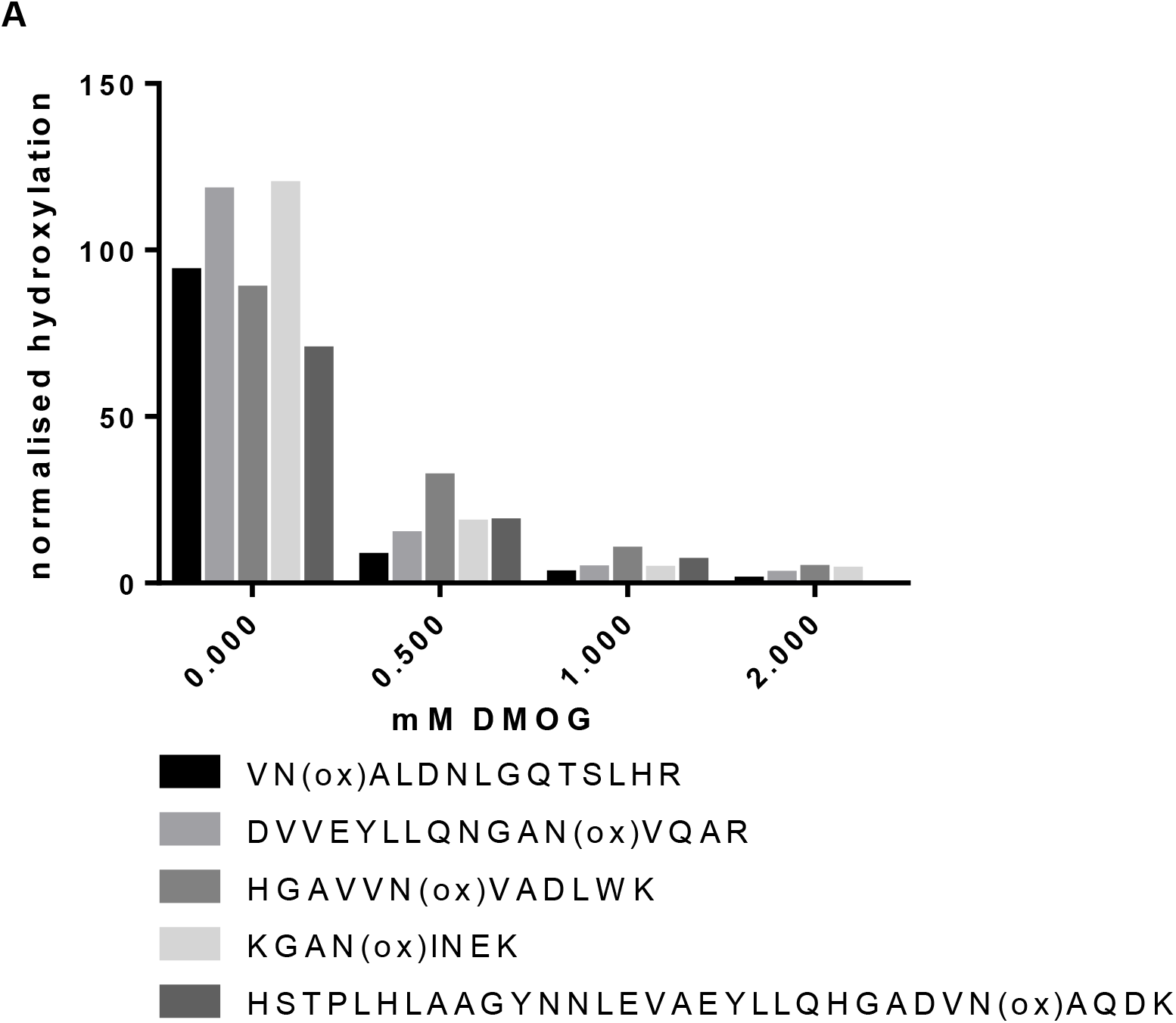
Determining the DMOG required to inhibit FIH. (A) HEK293T cells were transfected with FLAG-tagged TNKS2. 24hours post-transfection the media was replaced by heavy SILAC and cells were treated with different concentrations of DMOG or vehicle for four hours. FLAG-tagged proteins were immunoprecipitated, digested and analysed by mass spectrometry. Graph represents the intensity of heavy-labelled hydroxylated TNKS2 peptides in the presence of DMSO/DMOG, normalised by total heavy-labelled TNKS2 protein, n=2.

**Figure S2.**
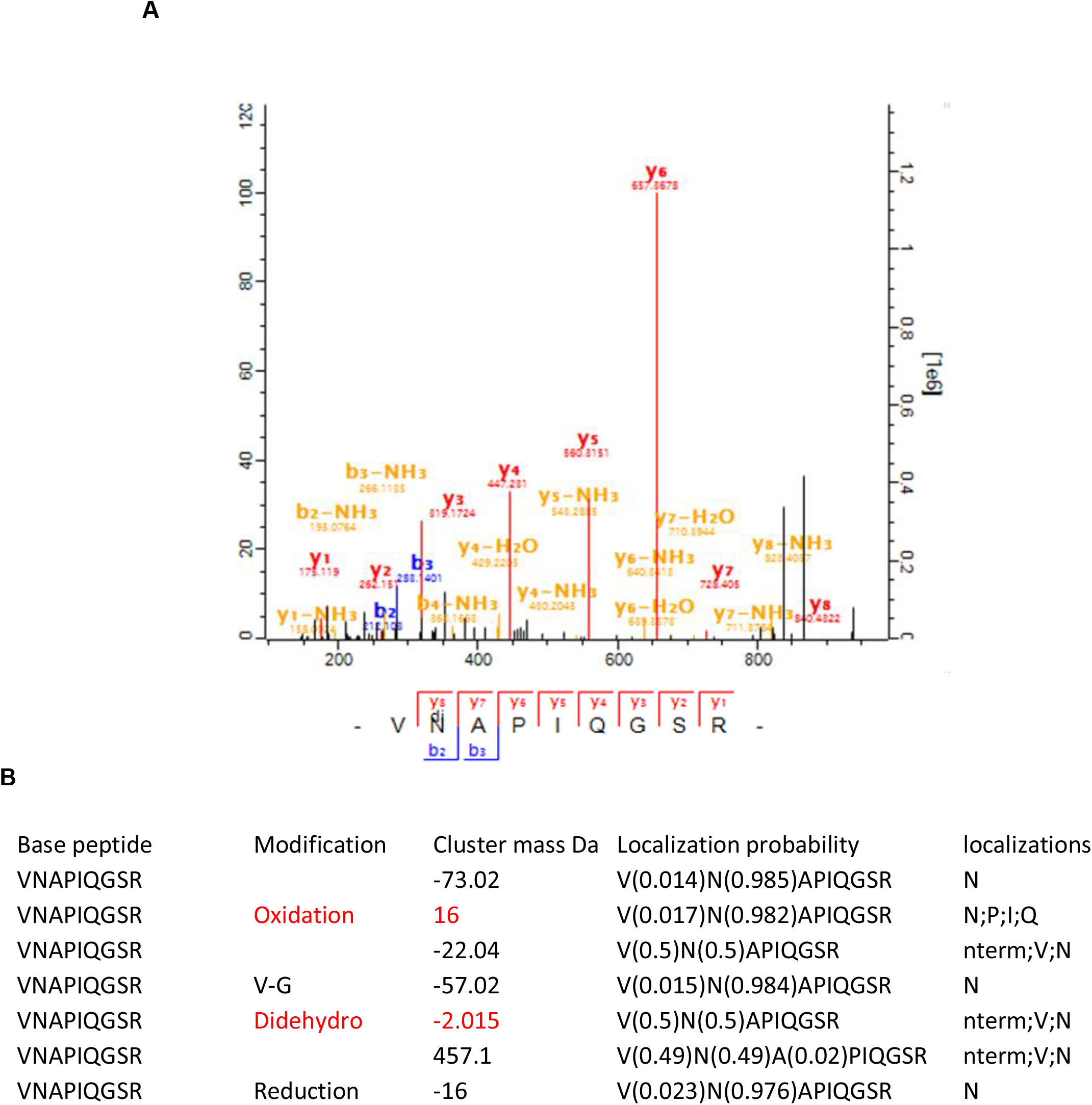
(A) MS/MS spectrum of the didehydrated HIF1α peptide (B) Table of dependent peptide modification identified matching VNAPIQGSR and localised on N

## References

Appelhoff, R.J., Tian, Y.M., Raval, R.R., Turley, H., Harris, A.L., Pugh, C.W., Ratcliffe, P.J., and Gleadle, J.M. (2004). Differential function of the prolyl hydroxylases PHD1, PHD2, and PHD3 in the regulation of hypoxia-inducible factor. J Biol Chem 279, 38458–38465.

Araos, J., Sleeman, J.P., and Garvalov, B.K. (2018). The role of hypoxic signalling in metastasis: towards translating knowledge of basic biology into novel anti-tumour strategies. Clin Exp Metastasis 35, 563–599.

Boisvert, F.M., Ahmad, Y., Gierlinski, M., Charriere, F., Lamont, D., Scott, M., Barton, G., and Lamond, A.I. (2012). A quantitative spatial proteomics analysis of proteome turnover in human cells. Mol Cell Proteomics 11, M111 011429.

Bruick, R.K., and McKnight, S.L. (2001). A conserved family of prolyl-4-hydroxylases that modify HIF. Science 294, 1337–1340.

Chan, D.A., Sutphin, P.D., Yen, S.E., and Giaccia, A.J. (2005). Coordinate regulation of the oxygen-dependent degradation domains of hypoxia-inducible factor 1 alpha. Mol Cell Biol 25, 6415–6426.

Chen, D.Y., Fabrizio, J.A., Wilkins, S.E., Dave, K.A., Gorman, J.J., Gleadle, J.M., Fleming, S.B., Peet, D.J., and Mercer, A.A. (2017). Ankyrin Repeat Proteins of Orf Virus Influence the Cellular Hypoxia Response Pathway. J Virol 91.

Ciechanover, A. (2005). Proteolysis: from the lysosome to ubiquitin and the proteasome. Nat Rev Mol Cell Biol 6, 79–87.

Cockman, M.E., Lippl, K., Tian, Y.M., Pegg, H.B., Figg, W.D.J., Abboud, M.I., Heilig, R., Fischer, R., Myllyharju, J., Schofield, C.J., et al. (2019). Lack of activity of recombinant HIF prolyl hydroxylases (PHDs) on reported non-HIF substrates. Elife 8.

Cockman, M.E., Webb, J.D., Kramer, H.B., Kessler, B.M., and Ratcliffe, P.J. (2009a). Proteomics-based identification of novel factor inhibiting hypoxia-inducible factor (FIH) substrates indicates widespread asparaginyl hydroxylation of ankyrin repeat domain-containing proteins. Mol Cell Proteomics 8, 535–546.

Cockman, M.E., Webb, J.D., and Ratcliffe, P.J. (2009b). FIH-dependent asparaginyl hydroxylation of ankyrin repeat domain-containing proteins. Ann N Y Acad Sci 1177, 9–18.

Duan, G., and Walther, D. (2015). The roles of post-translational modifications in the context of protein interaction networks. PLoS Comput Biol 11, e1004049.

Giaccia, A., Siim, B.G., and Johnson, R.S. (2003). HIF-1 as a target for drug development. Nat Rev Drug Discov 2, 803–811.

Gorres, K.L., and Raines, R.T. (2010). Prolyl 4-hydroxylase. Crit Rev Biochem Mol Biol 45, 106–124.

Gray, S.G., and Teh, B.T. (2001). Histone acetylation/deacetylation and cancer: an “open” and “shut” case? Curr Mol Med 1, 401–429.

Hunter, T. (2009). Tyrosine phosphorylation: thirty years and counting. Curr Opin Cell Biol 21, 140–146.

Hutton, J.J., Jr., Kaplan, A., and Udenfriend, S. (1967). Conversion of the amino acid sequence gly-pro-pro in protein to gly-pro-hyp by collagen proline hydroxylase. Arch Biochem Biophys 121, 384–391.

Ivan, M., Kondo, K., Yang, H., Kim, W., Valiando, J., Ohh, M., Salic, A., Asara, J.M., Lane, W.S., and Kaelin, W.G., Jr. (2001). HIFalpha targeted for VHL-mediated destruction by proline hydroxylation: implications for O2 sensing. Science 292, 464–468.

Jaakkola, P., Mole, D.R., Tian, Y.M., Wilson, M.I., Gielbert, J., Gaskell, S.J., von Kriegsheim, A., Hebestreit, H.F., Mukherji, M., Schofield, C.J., et al. (2001). Targeting of HIF-alpha to the von Hippel-Lindau ubiquitylation complex by O2-regulated prolyl hydroxylation. Science 292, 468–472.

Janke, K., Brockmeier, U., Kuhlmann, K., Eisenacher, M., Nolde, J., Meyer, H.E., Mairbaurl, H., and Metzen, E. (2013). Factor inhibiting HIF-1 (FIH-1) modulates protein interactions of apoptosis-stimulating p53 binding protein 2 (ASPP2). J Cell Sci 126, 2629–2640.

Karttunen, S., Duffield, M., Scrimgeour, N.R., Squires, L., Lim, W.L., Dallas, M.L., Scragg, J.L., Chicher, J., Dave, K.A., Whitelaw, M.L., et al. (2015). Oxygen-dependent hydroxylation by FIH regulates the TRPV3 ion channel. J Cell Sci 128, 225–231.

Kay, J.E., and Korner, A. (1966). Effect of cycloheximide on protein and ribonucleic acid synthesis in cultured human lymphocytes. Biochem J 100, 815–822.

Kiriakidis, S., Henze, A.T., Kruszynska-Ziaja, I., Skobridis, K., Theodorou, V., Paleolog, E.M., and Mazzone, M. (2015). Factor-inhibiting HIF-1 (FIH-1) is required for human vascular endothelial cell survival. FASEB J 29, 2814–2827.

Kivirikko, K.I., and Prockop, D.J. (1967). Hydroxylation of proline in synthetic polypeptides with purified protocollagen hydroxylase. J Biol Chem 242, 4007–4012.

Lando, D., Peet, D.J., Gorman, J.J., Whelan, D.A., Whitelaw, M.L., and Bruick, R.K. (2002a). FIH-1 is an asparaginyl hydroxylase enzyme that regulates the transcriptional activity of hypoxia-inducible factor. Genes Dev 16, 1466–1471.

Lando, D., Peet, D.J., Whelan, D.A., Gorman, J.J., and Whitelaw, M.L. (2002b). Asparagine hydroxylation of the HIF transactivation domain a hypoxic switch. Science 295, 858–861.

Larance, M., Ahmad, Y., Kirkwood, K.J., Ly, T., and Lamond, A.I. (2013). Global subcellular characterization of protein degradation using quantitative proteomics. Mol Cell Proteomics 12, 638–650.

Loenarz, C., and Schofield, C.J. (2011). Physiological and biochemical aspects of hydroxylations and demethylations catalyzed by human 2-oxoglutarate oxygenases. Trends Biochem Sci 36, 7–18.

Luo, W., Lin, B., Wang, Y., Zhong, J., O’Meally, R., Cole, R.N., Pandey, A., Levchenko, A., and Semenza, G.L. (2014). PHD3-mediated prolyl hydroxylation of nonmuscle actin impairs polymerization and cell motility. Mol Biol Cell 25, 2788–2796.

Mahon, P.C., Hirota, K., and Semenza, G.L. (2001). FIH-1: a novel protein that interacts with HIF-1alpha and VHL to mediate repression of HIF-1 transcriptional activity. Genes Dev 15, 2675–2686.

Mariotti, L., Templeton, C.M., Ranes, M., Paracuellos, P., Cronin, N., Beuron, F., Morris, E., and Guettler, S. (2016). Tankyrase Requires SAM Domain-Dependent Polymerization to Support Wnt-beta-Catenin Signaling. Mol Cell 63, 498–513.

Maxwell, P.H., Wiesener, M.S., Chang, G.W., Clifford, S.C., Vaux, E.C., Cockman, M.E., Wykoff, C.C., Pugh, C.W., Maher, E.R., and Ratcliffe, P.J. (1999). The tumour suppressor protein VHL targets hypoxia-inducible factors for oxygen-dependent proteolysis. Nature 399, 271–275.

McMahon, D. (1975). Cycloheximide is not a specific inhibitor of protein synthesis in vivo. Plant Physiol 55, 815–821.

Nguyen, L.K., Cavadas, M.A., Scholz, C.C., Fitzpatrick, S.F., Bruning, U., Cummins, E.P., Tambuwala, M.M., Manresa, M.C., Kholodenko, B.N., Taylor, C.T., et al. (2013). A dynamic model of the hypoxia-inducible factor 1alpha (HIF-1alpha) network. J Cell Sci 126, 1454–1463.

Ohsumi, Y. (2006). Protein turnover. IUBMB Life 58, 363–369.

Oka, O.B., and Bulleid, N.J. (2013). Forming disulfides in the endoplasmic reticulum. Biochim Biophys Acta 1833, 2425–2429.

Ploumakis, A., and Coleman, M.L. (2015). OH, the Places You’ll Go! Hydroxylation, Gene Expression, and Cancer. Mol Cell 58, 729–741.

Prass, K., Ruscher, K., Karsch, M., Isaev, N., Megow, D., Priller, J., Scharff, A., Dirnagl, U., and Meisel, A. (2002). Desferrioxamine induces delayed tolerance against cerebral ischemia in vivo and in vitro. J Cereb Blood Flow Metab 22, 520–525.

Ramchandani, S., Bhattacharya, S.K., Cervoni, N., and Szyf, M. (1999). DNA methylation is a reversible biological signal. Proc Natl Acad Sci U S A 96, 6107–6112.

Rehder, D.S., and Borges, C.R. (2010). Cysteine sulfenic acid as an intermediate in disulfide bond formation and nonenzymatic protein folding. Biochemistry 49, 7748–7755.

Rodriguez, J., Pilkington, R., Garcia Munoz, A., Nguyen, L.K., Rauch, N., Kennedy, S., Monsefi, N., Herrero, A., Taylor, C.T., and von Kriegsheim, A. (2016). Substrate-Trapped Interactors of PHD3 and FIH Cluster in Distinct Signaling Pathways. Cell Rep 14, 2745–2760.

Ruan, K., Song, G., and Ouyang, G. (2009). Role of hypoxia in the hallmarks of human cancer. J Cell Biochem 107, 1053–1062.

Schofield, C.J., and Ratcliffe, P.J. (2004). Oxygen sensing by HIF hydroxylases. Nat Rev Mol Cell Biol 5, 343–354.

Scholz, C.C., Rodriguez, J., Pickel, C., Burr, S., Fabrizio, J.A., Nolan, K.A., Spielmann, P., Cavadas, M.A., Crifo, B., Halligan, D.N., et al. (2016). FIH Regulates Cellular Metabolism through Hydroxylation of the Deubiquitinase OTUB1. PLoS Biol 14, e1002347.

Sharp, F.R., and Bernaudin, M. (2004). HIF1 and oxygen sensing in the brain. Nat Rev Neurosci 5, 437–448.

Singleton, R.S., Trudgian, D.C., Fischer, R., Kessler, B.M., Ratcliffe, P.J., and Cockman, M.E. (2011). Quantitative mass spectrometry reveals dynamics of factor-inhibiting hypoxia-inducible factor-catalyzed hydroxylation. J Biol Chem 286, 33784–33794.

Snell, C.E., Turley, H., McIntyre, A., Li, D., Masiero, M., Schofield, C.J., Gatter, K.C., Harris, A.L., and Pezzella, F. (2014). Proline-hydroxylated hypoxia-inducible factor 1alpha (HIF-1alpha) upregulation in human tumours. PLoS One 9, e88955.

Tonks, N.K. (2006). Protein tyrosine phosphatases: from genes, to function, to disease. Nat Rev Mol Cell Biol 7, 833–846.

Trewick, S.C., McLaughlin, P.J., and Allshire, R.C. (2005). Methylation: lost in hydroxylation? EMBO Rep 6, 315–320.

Waaler, J., Machon, O., Tumova, L., Dinh, H., Korinek, V., Wilson, S.R., Paulsen, J.E., Pedersen, N.M., Eide, T.J., Machonova, O., et al. (2012). A novel tankyrase inhibitor decreases canonical Wnt signaling in colon carcinoma cells and reduces tumor growth in conditional APC mutant mice. Cancer Res 72, 2822–2832.

Walsh, C.T., Garneau-Tsodikova, S., and Gatto, G.J., Jr. (2005). Protein posttranslational modifications: the chemistry of proteome diversifications. Angew Chem Int Ed Engl 44, 7342–7372.

Yang, E., Tacchelly-Benites, O., Wang, Z., Randall, M.P., Tian, A., Benchabane, H., Freemantle, S., Pikielny, C., Tolwinski, N.S., Lee, E., et al. (2016). Wnt pathway activation by ADP-ribosylation. Nat Commun 7, 11430.

Yu, F., White, S.B., Zhao, Q., and Lee, F.S. (2001). HIF-1alpha binding to VHL is regulated by stimulus-sensitive proline hydroxylation. Proc Natl Acad Sci U S A 98, 9630–9635.

